# Fungal phylogeny reconstructed using heterogeneous models reveals new placement of Microsporidia

**DOI:** 10.1101/2023.06.28.546859

**Authors:** Ignacio Riquelme-Medina, James O. McInerney

**Author notes:** To whom correspondence should be addressed. Email: Ignacio Riquelme-Medina James O. McInerney.

## Abstract

Fungi have significant impacts on global ecosystems, playing roles in organic matter decomposition, as disease agents, as well as having both positive and negative economic effects. Fungal evolutionary history remains partly unresolved with the presence on many *incertae sedis* clades, lacking a robust placement on the fungal phylogenetic tree. Some of the more problematic groups whose placement remains unclear are the intracellular parasites *Microsporidia* and Cryptomycota, both of whom have accelerated rates of nucleotide substitution and reduced genomes, which makes the resolution of these groups hard. In this project we used tree and data heterogeneous models, that can account for different substitution rates between different protein families as well as different parts of the phylogenetic tree, to reconstruct the phylogeny. We recovered a well-supported topology for much of the fungal phylogeny as well as a robust placement for both *Microsporidia* and Cryptomycota, the former being rooted deeply within the fungi and the latter a placement on a sister branch to the fungi.

## Introduction

Fungi make a significant impact on planetary ecosystems (1), and on human activity including in the food and beverage industries, acting as pathogens in plant and animal diseases, and in research, serving as model organisms, and assisting in drug discovery. Fungi are characterized by being highly variable eukaryotes, with simple morphologies (typically filamentous but they can also be unicellular), sharing a common ancestor with animals dating to approximately 1 billion years ago (2). Fungi have relatively small genomes and are heterotrophic (3), they can have haploid and diploid stages in their life cycle and do not have movement except for some flagellated spores. They can be saprobes, symbionts and parasites of plants, animals, or other fungi, but they are not able to photosynthesise. Owing to their variability they have adapted repeatedly to the same kind of environments at different points in their evolutionary history and the emergence of divergent phenotypes such as parasitism and mutualism, are not exclusively found in a single clade (3). As an example of how quickly fungi can adapt, we can look at the Hymenochaetales order, which has saprobes, mycorrhiza, and strong and weak plant parasites amongst its members (4). A fungal phylogenetic tree has been recently constructed by the Assembling the Fungal Tree of Life project (AFToL) (5, 6). However, many branches of the phylogeny are not well resolved, resulting in organisms that are classified as i*ncertae sedis* as they appear in the NCBI database taxonomy (7). Some of the deeper branches, for example *Microsporidia* and *Cryptomycota* (*Rozella*), also have an uncertain phylogenetic placement, and it is unclear whether they belong to the fungal kingdom or are sister groups to the fungi (8–14).

Fungi have long been good model organisms for eukaryotes, owing to their small genome size, the fact that they are easy and inexpensive to maintain, they are relatively amenable to genetic manipulation, and that some form tissues that allow transcriptomic experiments (15–19). In recent years efforts to sequence complete fungal genomes have intensified, particularly with the 1,000 Fungal Genome Project (FGP) (20) which is a project to sequence 1,000 fungal genomes and make them accessible to the public. With the FGP hundreds of fungal genomes have already been sequenced but remain to be analysed. Here we report the use of these data to construct an updated phylogeny of the fungi, paying special attention to unresolved groups.

There is ongoing discussion about the inclusion of *Microsporidia* and *Cryptomycota* in the fungal kingdom (21). The difficulty associated with placing these groups on the eukaryote phylogeny is due to their specialised obligate intracellular parasitic lifestyle. This lifestyle has led to considerable genome reduction, and unusually high nucleotide and amino acid substitution rates, as well as the loss of their mitochondria (22–24). High substitution rates are known to be problematic for accurate phylogeny reconstruction (25, 26), and several solutions have been proposed to mitigate these effects (27–29). In this paper we tested whether using a model that specified a homogenous evolutionary process would accurately capture the signals in the data. We found that such a model was a poor fit to the data and to properly account for the signals in the data we used both tree-heterogeneous and sequence-heterogeneous phylogenetic reconstruction approaches (30, 31). A compositionally heterogeneous model specifies more than one evolutionary process for the different genes or proteins used for a supermatrix reconstruction as well as being able to use several models of evolution for different branches of a phylogeny. By allowing the phylogenetic reconstruction to account for different rates of evolution for different genes and branches we should be able to account for the genome reductions observed in *Microsporidia* and *Cryptomycota*.

## Methods

### Dataset

A total of 690 genomes available through the Fungal Genomes Project (FGP) (20) were downloaded using the Globus data sharing tool (32). The DNA sequences, protein sequences, annotations, and functional annotations were obtained, using the filtered models for each genome. These files are included for every genome in the FGP database, and include GO (33), KEGG pathways (34), InterPro (35), KOG (36), and SignalP (37) annotations.

To evaluate the quality of the genomes we used the Fungal Genome Mapping Project (FGMP) (38) a framework to check for the presence of several conserved and ultraconserved genes. Poor quality genomes were identified as having fewer than 75% of the conserved genes present and were discarded leaving a total of 671 genomes. Owing to their known reduce genome size, parasitic fungi were not discarded at this step despite possessing less than 30% of conserved genes.

To deal with the size of our dataset and its associated metadata a SQL database was built using PostgreSQL versions 11 to 12.2. In this database we built a protein table where the protein identification, protein sequence and protein family of each protein was stored. Other tables were built to store the functional annotations and were connected to the proteins by the identification numbers provided by FGP. Finally, a table with all the fungal species and their categories (WoL, habitat, infectious, extremophile and notes) was connected to the protein families in the protein table by the edge information obtained in the previous step with MCL. The structure of the database can be seen in Supplementary Figure 1. The python package psycopg2 (https://pypi.org/project/psycopg2/) was used to interact and retrieve information from the database.

### Data Processing

Protein sequences were then used to perform an all-versus-all BLAST (39, 40) search (BLASTP v2.4.0, e value = 1*e-6). The BLAST output was processed using the Markov Cluster Algorithm (MCL) (41) version 14-137 to detect clusters. To select an appropriate inflation value for our dataset we used the highest inflation value where most conserved proteins (*e.g.* ribosomal proteins) were recovered in the same cluster. The final inflation value was set to 1.4. This resulted in the recovery of 965,828 clusters from the dataset.

We constructed phylogenetic hypotheses using a concatenation of several alignments. Therefore, we sought to identify clusters that include almost all 671 taxa, where the clusters contained little, to no, duplicated genes. We identified an initial set of 507 clusters that had 671 ± 20 taxa, allowing for some gene duplications, and that included at least 95% of all the taxa to avoid clusters with high number of duplications.

### Phylogenetic Analyses

Multiple sequence alignment was carried out using MAFFT v6.611b (42) (auto1 option) as an alignment tool. We then used Trimal v1.2 (43) (automated1 option) to remove poorly aligned positions in the alignments. Prottest 3 (44) was used to assess the best substitution model for each of the alignments. After assigning the best model to each alignment, phylogenetic hypotheses were constructed using RAxML v8 (45). Since gene duplications have been shown to be a source of discrepancies between a gene tree and a species tree, often because duplicated genes evolve at different rates (46), we tested further the resulting 507 trees to check whether gene duplications were few and evolutionarily close in each tree. If the duplications are evolutionarily close, it would mean that there were few nucleotide substitutions between duplicated genes and that the different rate of evolution would have less impact on the gene tree. This was checked using ETE3 (47) node distance comparison function in python v2.7. To do this a custom script was made that checks the distance between all the duplicated gene pairs in one tree and reports the gene tree as valid to construct a species tree if: a) the duplicates are in very close proximity (same branch) and b) if no more than one pair of duplicates is further away than this set distance. Using this relaxed criterion, a total of 58 gene trees were selected. In addition, out-groups were added to each gene unaligned file, using BLASTP as described before to search for similar proteins in 4 different organisms: a mammal (*Homo sapiens*), a cnidarian (*Nematostella vectensis*), a choanoflagellate (*Monosiga brevicollis*) and a plant (*Arabidopsis thaliana*) making a total of 675 taxa in the dataset. The gene sequences were aligned again with the outgroups included as previously described and then the aligned gene sequences were concatenated to construct the species tree. If one gene tree was missing a taxon from the complete species tree, the gap was filled with missing characters of the same length as the gene.

The resulting concatenated alignment was of 11,559 amino acids in length. The programs Prottest 3 and RAxML were used as described previously to construct several phylogenetic hypotheses using a single model for all parts of the alignment. Bootstrap resampling (48) using 100 replicates was used to assess support for internal branches. The same alignment file was used with a heterogeneous model tree using the P4 software program (30). P4 implements both data-heterogeneous and tree-heterogeneous models using a modified version of MCMC. Since heterogeneous models are much more computationally expensive than homogeneous models, the original alignment file could not be used to construct a heterogeneous tree in a reasonable amount of time. Therefore, we had to make a reduced dataset using a selection of taxa from the original alignment. We chose a single taxon to represent each uncontroversial group in the maximum-likelihood tree apart from the out-group. However, the *Microsporidia* branch and *Rozella* were left intact as they were the most problematic branches in the original tree. The number of taxa in the reduced alignment was 59.

First to account for the data heterogeneity, the alignment file was used as input for the PartitionFinder 2.11 program (49). PartitionFinder searches for differing rates of substitution across regions described by the user in an alignment with the purpose of dividing them in partitions of proteins or genes with similar evolutionary rates and assigning a phylogenetic model to each partition. In our case we split the alignment in to each one of the partitions indicated by PartitionFinder (Partitions = 27). Afterwards the partition data and the alignment were analysed using P4, assigning a model to each partition in accordance with PartitionFinder results. Furthermore, each partition was also assigned the second-best model found by PartitionFinder to account for heterogeneity across the tree. Finally, MCMC was run for 300,000 generations, assessing whether the MCMC chains had converged at the end of the process. Additionally, the tail area probability of the MCMC run was calculated using a function included in P4 to assess whether the phylogeny recovered by MCMC could plausibly explain our original data.

The final step in the analysis was to combine the results obtained from the heterogeneous model tree to the full dataset. To do this we used PartitionFinder again with the full dataset since RAxML allows for data-heterogeneous models, obtaining a total of 51 partitions. Each partition was assigned the substitution model indicated by PartitionFinder for RAxML. To avoid problems with long branch attraction in this tree hypothesis we forced RAxML to root the tree on the out-group, as it was supported by the reduced dataset phylogenetic tree. After the phylogenetic analysis was finished, bootstrap resampling with 100 replicates was used to assess support for internal branches.

### Phylogenetic Tree Comparisons

To understand the similarity of our phylogenetic trees to other published phylogenies, we compared the complete dataset phylogenetic tree with other phylogenies using the program TOPD/FMTS (50). The phylogenies were obtained and downloaded from the TreeBASE database (51, 52), selecting phylogenies that included taxa from our dataset. This software allows to calculate the nodal distance and the split distance, also known as the Robinson–Foulds metric (53), between any given pair of phylogenetic trees. To draw the tanglegrams between pairs of phylogenetic trees we first used the R package Phylogram v2.1.0 (54) for converting phylograms into dendrograms, and then we used the R package Dendextend v1.17.1 (55) to create the tanglegrams from the resulting pairs of dendrograms in R v4.2.3 (56).

We also compared the topologies we recovered to a well-supported fungal phylogeny proposed by McCarthy and Fitzpatrick (57), that uses a similar but smaller dataset from the FGP. This comparator dataset does not contain the more problematic groups in the fungi, like *Microsporidia*, which would remove any artefact introduced by these groups in the phylogeny.

We first checked whether the species present in the McCarthy and Fitzpatrick dataset were present in our dataset, having passed our genome quality evaluation step. For species in the McCarthy and Fitzpatrick dataset that were not present in our dataset, but had other close species in the same family, we used this species instead. Specifically, *Zymoseptoria tritici* was changed for *Zymoseptoria ardabiliae, Candida albicans* for *Candida tanzawaensis, Microbotryum lychnidis-dioicae* for *Microbotryum violaceum* and *Rhizopus oryzae* for *Rhizopus microsporus.* Species that did not appear in our dataset and did not have a close relative were excluded from the analysis, specifically *Endocarpon pusillum, Orpinomyces sp. C1A,* and *Batrachochytrium dendrobatidis.* Instead of using an external source genome for *Allomyces macrogynus* that was used in the paper dataset, we used a genome from the FGP instead. This resulted in a total of 81 species that were used in this comparison step for the phylogenetic analysis. This dataset will be referred to as comparison dataset from this point onwards.

We then extracted a subset from our BLAST output that included only edges between the species present in the validation dataset and proceeded to apply the same steps as with our phylogenetic analysis. We used MCL with the same inflation value, 1.4, to obtain a total of 155,969 protein clusters. Afterwards, we filtered the clusters to select candidate genes for the phylogenetic analysis with a stricter filter due to the high number of genes returned if we applied the same criteria as with our dataset. We filtered only for clusters that had every species in the dataset present and at most two duplications, which left a total of 35 candidate clusters. Using the same procedure as with our dataset, we selected a total of 21 clusters that had either no duplications or duplications near one another. For clusters that had duplications, one of them was removed randomly to leave only one protein per species. As we did with the analysis of our new dataset, the selected clusters were aligned, trimmed, and concatenated with the same settings as with the original dataset to form a concatenated alignment with a final length of 7,611 amino acids. We used PartitionFinder again to determine the heterogeneity of substitution rates in the data (Validation Partitions = 13). Finally, we used two different P4 analyses, one considering only data-heterogeneity, which included the model indicated by PartitionFinder for each partition, and one considering data and tree-heterogeneity, which used two models indicated by PartitionFinder for each partition. Both analyses MCMC processes were executed until convergence was achieved, using 100,000 generations for the data-heterogeneous analysis and 240,000 generations for the data and tree-heterogeneous analysis. The tail-area probability of each analysis was also calculated using P4.

### Supertree

We also investigated phylogeny reconstruction of our dataset using a supertree approach. We used Clann (58), which implements several methods of supertree construction. We focussed on the Matrix Representation with Parsimony (MRP) method. MRP uses multiple individual gene trees to collect evidence for branch split support. Branch splits are extracted from phylogenetic trees and stored in the form of a matrix of ones and zeros. Using the maximum parsimony criterion, along with a tree search method, the matrix is used to construct the final supertree that minimises the number of character state changes in the dataset.

Thus, we used the clusters obtained by applying MCL to the complete dataset, with each cluster being subsequently treated as a gene family, to construct the individual gene trees. We filtered the clusters to only include clusters with no duplications, since gene duplications introduce problems that are difficult to deal with using current supertree methods (59). We also only used clusters that contained four taxa or more. This approach resulted in a total of 49,261 gene families being used for supertree construction. Each gene family was then aligned and trimmed in the same way as with the supermatrix dataset (MAFFT auto1 option and Trimal automated1 option). Afterwards, Prottest 3 was used to assign the appropriate substitution model to each gene alignment and then gene trees were constructed using RAxML with the indicated substitution model for each tree. Finally, we used Clann to construct the supertree by using the MRP method with default parameters for heuristic tree search (Parsimony analysis and tree bisection and reconnection as the type of heuristic search) with no repetitions.

## Results

### Phylogenetic Analysis

As an initial step in our phylogenetic analysis, we estimated several phylogenetic hypotheses under the LG+I+G model in RAxML with bootstrap resampling support for the total amount of taxa in the dataset. These trees had 675 leaves (671 fungi and 4 out-group taxa). The hypotheses recovered the phylum and class groups in the fungi with high confidence, particularly in the *Dikarya*. However, there was no identifiable split between the in-groups and the out-groups. Moreover, in some hypotheses *Microsporidia* appear branching amongst the out-group taxa, an example of which can be seen in Figure ***1***. Deep branches in hypotheses where the out-group appears split showed low support values. So, while the maximum-likelihood tree has provided good results for resolving short fungal branches, it failed to resolve the deep branches of the tree, presumably due to elevated substitution rates are within the *Microsporidia* clade, compared with the rest of the tree. This difference in branch length suggests very different rates of evolution, and previously it has been shown that *Microsporidia* have many fast-evolving genes (60), when compared to the rest of the fungi.

**Figure 1:**
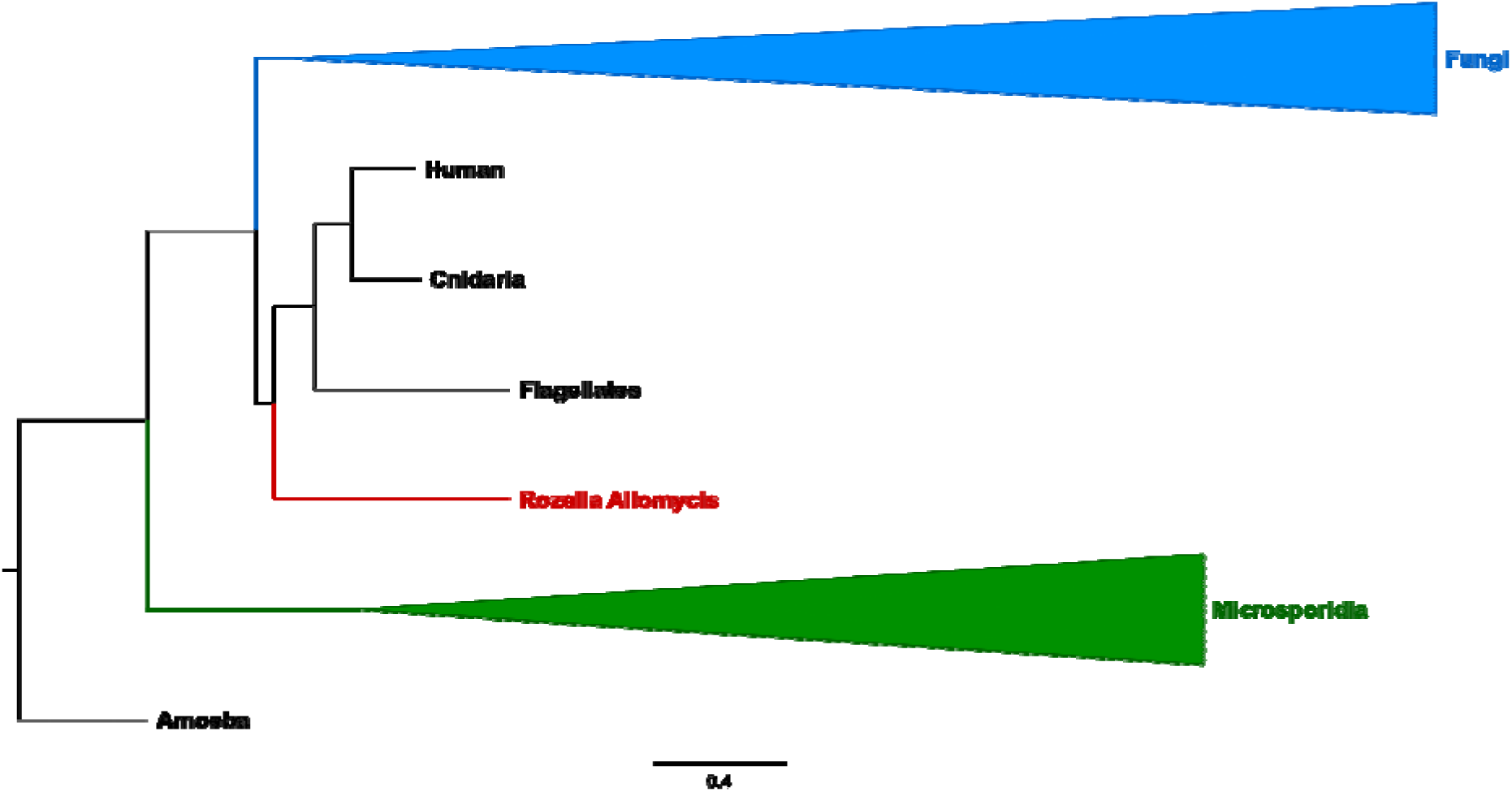
Example of a problematic phylogeny where the out-groups appear as a non-monophyletic clade, with *Microsporidia* and *Rozella* appearing within the outgroups.

To overcome problems associated with low support for the placement of long branches on our tree, we used heterogeneous models, which can account for variation in rates of evolution across different proteins, different sites within proteins and different branches across the tree. Because heterogeneous models require much more computational resources compared with homogeneous models, we used a reduced dataset that consisted of a representative taxon for each clade in the fungal in-group, though we retained all out-group sequences and *Microsporidia*.

Phylogenetic hypotheses were constructed from the reduced dataset using P4 using two models per partition to account for tree-heterogeneity of substitution rates using the substitution models indicated by PartitionFinder. This resulted in two LG+I+G substitution models per partition, for twenty-four partitions, and two LG+G substitution models per partition for the remaining three partitions (LogLikelihood = -557,634.03). From the 300,000 generations of the MCMC run, the first 200,000 were discarded as burn-in, and a majority-rule consensus tree was constructed from the remaining 100,000 generations. Branch split support values were obtained from the posterior probability distribution after MCMC had converged. According to these data and this model, Microsporidia can be placed as a sister group to Dikarya with strong support, and Rozella can be placed outside the Fungi as can be seen in Figure ***2***. There is a complete split between the in-groups and the out-groups of the tree unlike the previous tree, which also indicates that when accounting for different rates of evolution with the heterogeneous models helps resolve deep branches.

**Figure 2:**
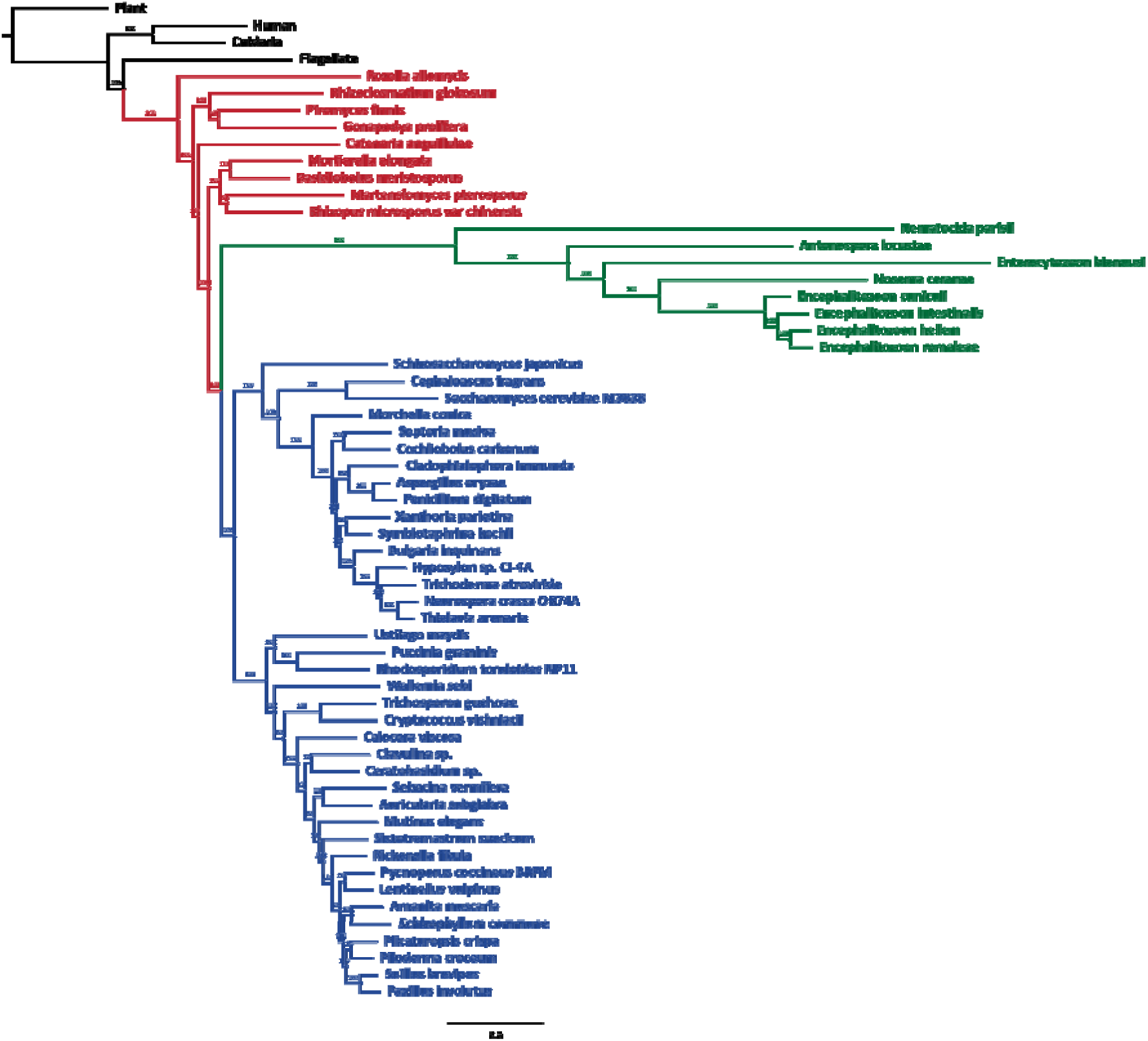
Reduced dataset data and tree heterogeneous Phylogenetic tree. Each branch is marked with the Posterior Probability Distribution (PPD) percentage of support. The out-groups are in black, early fungi are coloured red, *Microsporidia* are coloured green and Dikarya are coloured blue.

Two further MCMC runs were carried out to check whether they recovered phylogenies with similar topologies. Split support standard deviation between MCMC processes was measured at 0.0719. We also examined the tail area probability of the phylogenetic hypothesis using Posterior Predictive Simulations and this simulation did not reject the NULL hypothesis that the real data was no different to the simulated data under this model (p = 0.1984), which means that our dataset could have plausibly been generated by the model. Finally, we checked the extent to which the likelihood of the phylogenetic hypotheses was improved by using a data-and-tree heterogeneous model, so we ran another two MCMC processes, one with a data-heterogeneous model, and another with a homogeneous model, for comparison. For the data-heterogeneous model, we used a single substitution model per partition using the substitution models indicated above, and the MCMC chains were run for 300,000 generations. This resulted in a substantially lower LogLikelihood score of -559,060.90, with a tail area probability, p = 0.1236. This data-heterogeneous model topology is more like other topologies in the literature, placing Microsporidia as a sister group to the Fungi together with Rozella. For the homogeneous model we used a single LG+I+G model and run the MCMC for 300,000 generations. As expected, this resulted in an even lower Log Likelihood score of -562,923.74, with a tail area probability p = 0.409). The homogeneous tree topology is like the data-heterogeneous topology, with Microsporidia being placed outside of the Fungi together with Rozella. However, even if the homogeneous model can plausibly explain the data, some *Microsporidian* taxa appear spread across the tree within the fungi with low Posterior Probability support, and the likelihood of the model is much worse when compared with the more complex models. Next, we compared the different heterogeneous likelihood models by carrying out a likelihood ratio test. According to the test of the significance of the difference in log-likelihood scores between the trees, the data-and-tree heterogeneous model is a significantly better fit to the data than the data heterogeneous model (*P* value < 0.001). In summary, simulations suggest that both the data-and-tree heterogeneous hypothesis and the data heterogeneous hypothesis with their respective topologies are plausible hypothesis for a fungal phylogeny, but the data-and-tree heterogeneous model is a significantly better hypothesis according to the likelihood of the model.

Finally, we defined partitions again with the complete alignment using PartitionFinder to account for varying rates of amino-acid substitution between proteins in the maximum-likelihood hypothesis with the complete dataset. Then we used the root that we obtained with the reduced dataset heterogeneous model we built a final ML tree defining the out-groups as indicated by the heterogeneous model’s hypothesis. This last phylogenetic hypothesis was constructed using RAxML under the LG4X+I+G substitution model with the data split into thirty partitions, the LG4M+I+G substitution model for sixteen partitions, LGF+I+G substitution model for two partitions and LG+I+G for the remaining two partitions as indicated by PartitionFinder. Then we assessed support for the tree internal branches of this tree by using bootstrapping resampling (100 pseudoreplicates) as shown in Figure 3. The branches of this tree were collapsed in clades matching the NCBI’s fungal taxonomy database to allow for a better visualization.

**Figure 3:**
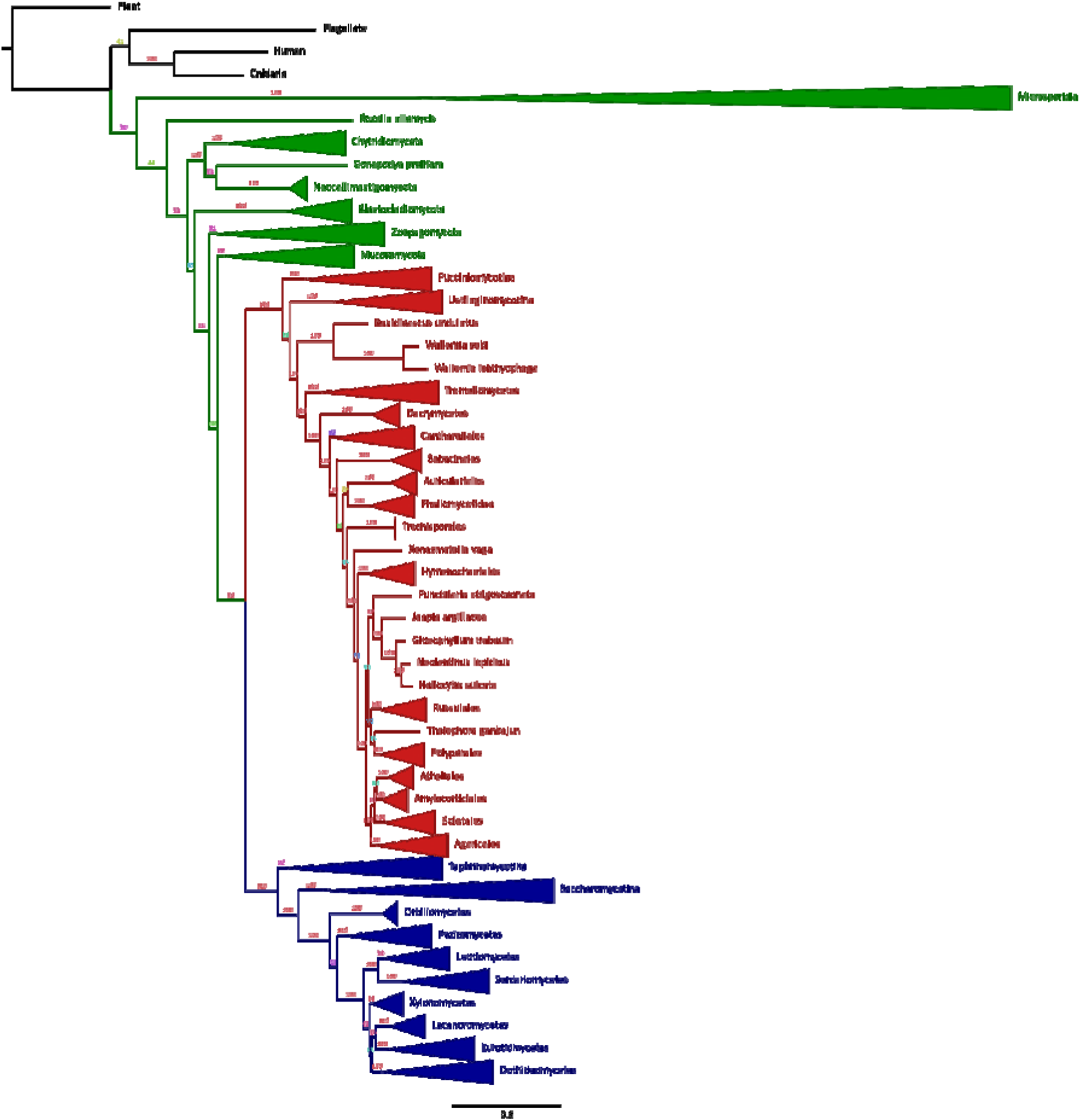
Fungal phylogenetic tree with bootstrap values. In Green are the groups outside of Dikarya, Red is Basidiomycota, Blue is Ascomycota and Black is the outgroup.

**Figure 4:**
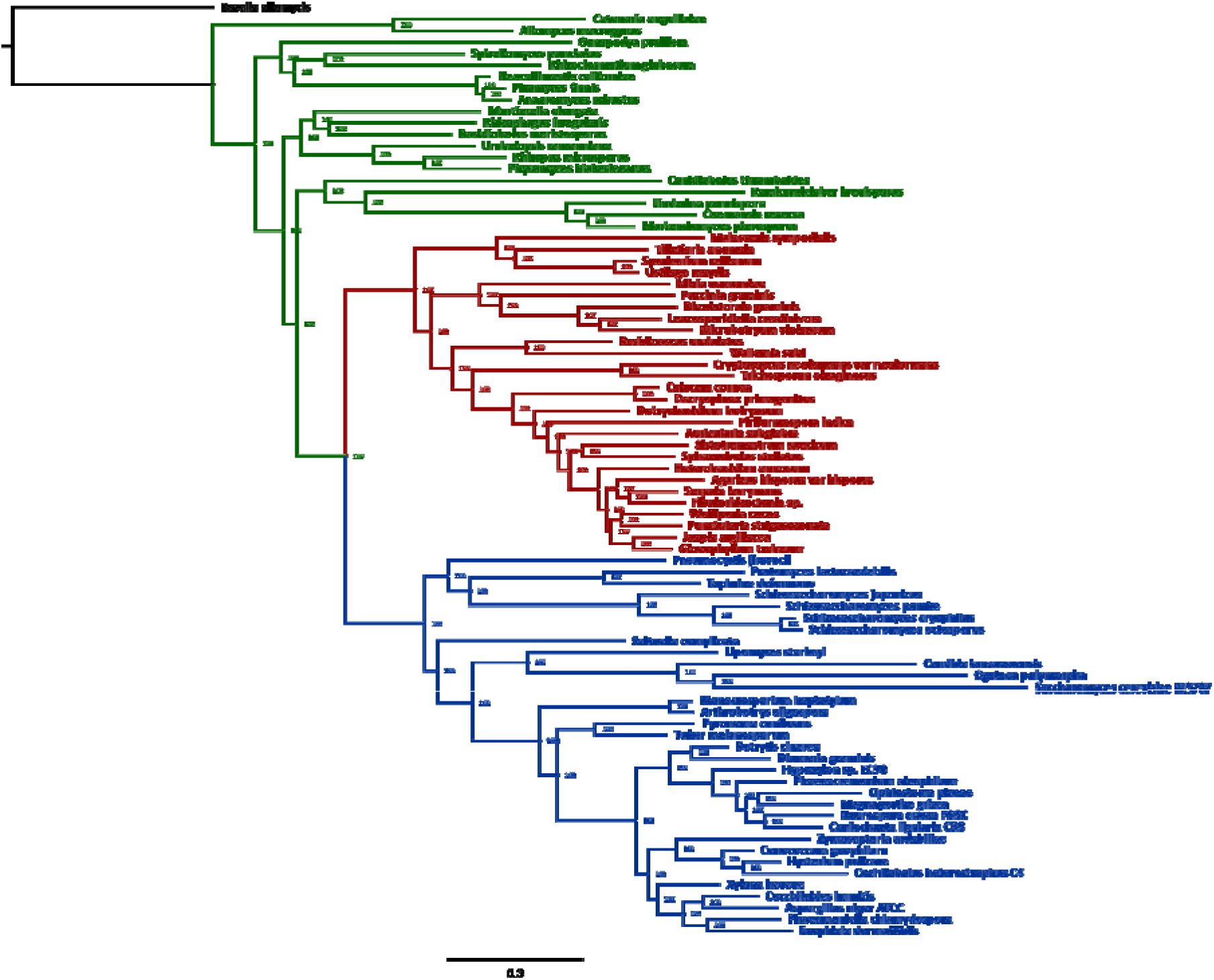
Validation dataset data and tree heterogeneous Phylogenetic tree. Each branch is marked with the PPD percentage of support. In Green are the early fungi, Red is Basidiomycota, Blue is Ascomycota and Black is the outgroup.

The resulting trees obtained from the concatenated analysis are very similar to the current taxonomy of the fungi according to NCBI. Since the leaves in the tree in Figure 3 were collapsed according to NCBI’s taxonomy many of the groups found by our analysis are in agreement with the current consensus. This tree will be referred as the Final Tree and will be the main basis of our discussion.

### Phylogenetic comparisons

Finally, we have compared our complete phylogeny with several previously recovered phylogenies from other studies. The results obtained after analysing each pair of trees with TOPD/FMTS, *i.e.,* our complete phylogeny compared to each of the literature phylogenies by using TOPD/FMTS nodal distance and split distance modes, can be seen in Table 1.

As for the alternative supertree method, the phylogeny that can be seen in Figure **Error! Reference source not found.** was obtained by using the MRP method with a total of 49,261 single gene trees. Bootstrap analysis of the data could not be performed due to the long time it would take to complete for 100 iterations, as each iteration took several weeks to complete.

**Table 1:**
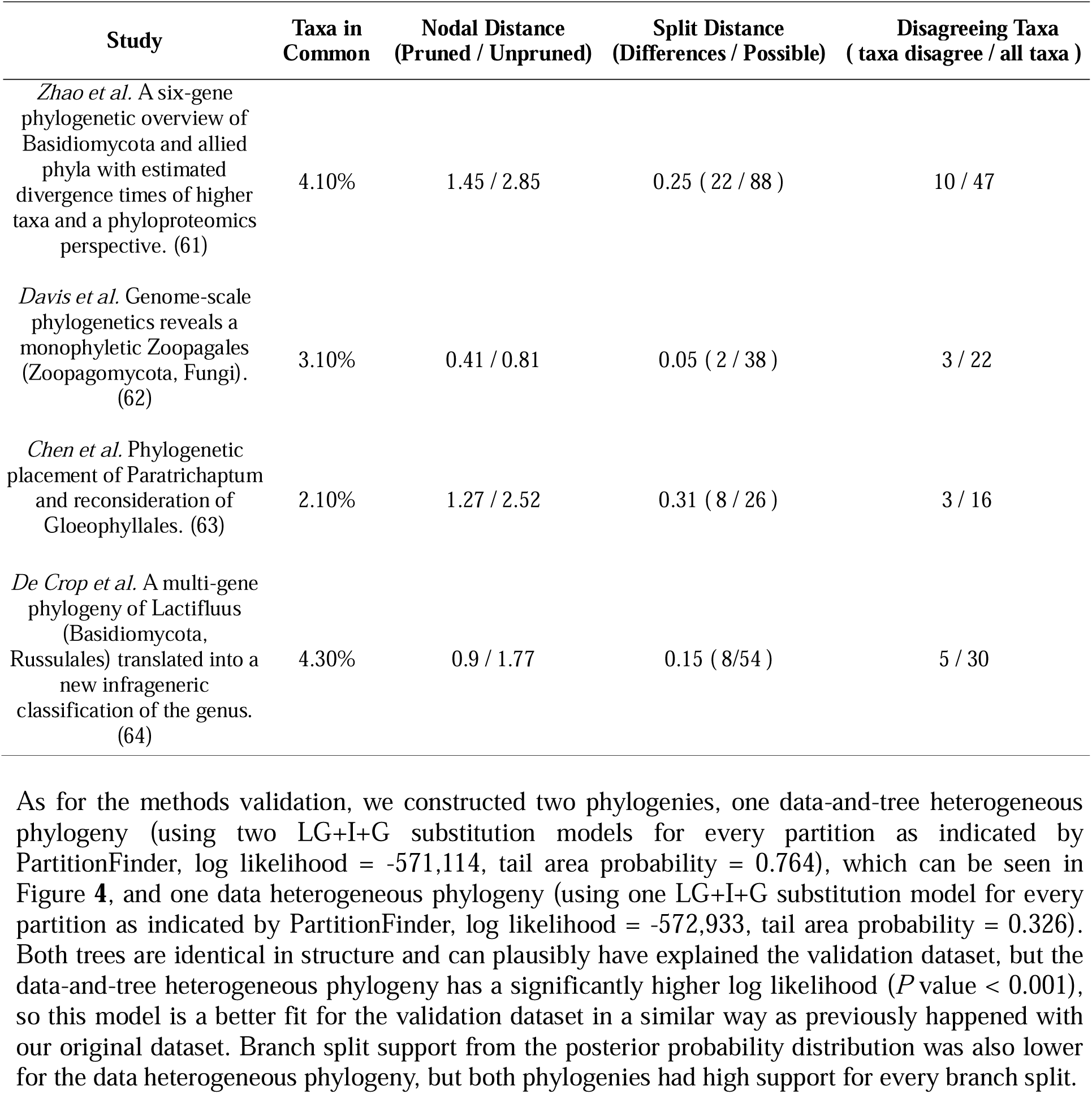
TOPD/FMTS results of the comparison between the recovered phylogeny of the total taxa with the phylogeny proposed by each one of the following studies.

## Discussion

The supertree approach was incapable of properly separating the two big clades present in Dikarya, Ascomycota and Basidiomycota. While in both our supermatrix approach and in the literature these groups are easily recovered, in the supertree approach that we implemented, Ascomycota and Basidiomycota are mixed in some branches. For example, in Figure **Error! Reference source not found.** Taphrinomycotina (Ascomycota) appears as a sister branch of the Tremellomycetes (Basidiomycota) or Saccharomycotina (Ascomycota) as a sister branch of Wallemiomycetes (Basidiomycota). There are also issues with the placement of the fungal groups outside of Dikarya, where clades like Zoopagomycota are not monophyletic and are instead separated and mixed with other groups like Mucoromycota, or even appear to branch within the Dikarya phylum, for example, *Ramicandelaber brevisporus*, is placed within Pucciniomycotina. Other problematic groups like Microsporidia appear mostly within Dikarya but are also not a monophyletic group and are spread across several branches. In contrast, the supertree approach recovers well-established clades that have a more recent common ancestry, and many of the smaller groups on the phylogeny are in accordance with NCBI’s taxonomy and appear as collapsed clades in Figure **Error! Reference source not found.**.

One possible explanation for why the supertree phylogeny struggled to reconstruct the deeper branches of the phylogeny can be the distribution of the gene tree size of our dataset. Almost 90% of the single gene trees used for the supertree analysis have 7 taxa or fewer, which is 1% of the total species in our dataset. This kind of data may provide enough resolution to resolve smaller clades but not enough to resolve the deep branch splits in the phylogeny. The size of our dataset might be the cause of this issue as it might make the presence of single gene trees without duplication sparse. We faced the same issue previously when we were selecting the protein families to use with our supermatrix approach. For our large supermatrix approach, we needed to set lax filters and allow for some duplication events to find enough candidate genes. On the other hand, with the smaller validation supermatrix the filters to select candidate proteins needed to be stricter due to the large number of candidate protein families retrieved by the previous filters. Therefore, the large size of our dataset can explain the lack of suitable protein families needed for supertree reconstruction. The issues with problematic groups in the supertree, namely that some of them are not monophyletic, might be explained by the presence of intracellular parasites like in Microsporidia, which have greatly reduced genomes and have even lost or repurposed ribosomal proteins (65, 66), which implies that single gene trees where these species are present are much sparser. Other possible cause for the low resolution of the supertree is that single gene trees may have a similar problem as the supermatrix approach, where homogeneous substitution models are enough to model the evolutionary process in groups with recent common ancestry, but the more complex heterogeneous models are needed to model the evolutionary process of deep branches and problematic groups. These issues indicate that supertree phylogeny reconstruction may not be a good fit for our particular dataset.

In contrast to the supertree approach, the supermatrix approach could recover uncontroversial splits between large fungal clades, robustly splitting Dikarya from early fungi and Ascomycota from Basidiomycota, even when using only homogeneous models. Additionally, due to the size of our dataset, the supertree methodology involved several weeks of computation time to construct a single supertree, which also hampered the calculation of branch split support through bootstrap analysis. Given the low resolution of the supertree approach, the long computational time involved, and the difficulty in procuring bootstrap support values for branch splits, we decided to focus our efforts on using tree and data heterogeneous models with the supermatrix approach.

We compared our tree with the topology from Ebersberger et al (67). Even though our tree has many more taxa, both phylogenetic trees share a similar structure when it comes to support for large clades. There are some minor differences among the Basidiomycota like *Gloeophyllum* appearing in a deeper branch than *Heterobasidion* in our tree and that *Schizophyllum* and *Pleurotus* are in different clades in our tree. However, these differences seem to be due to varying number of taxa in these positions of the tree between the two phylogenies. In the case of the Ascomycota, the clade of interest in our tree is somewhat more densely populated than on Ebersberger’s phylogeny. Conversely, in the case of the Basidiomycota, there are considerably more taxa in Ebersberger’s phylogeny than in ours. Consequently, the difference between number of taxa in these branches between phylogenies may play a role in explaining their differing topologies. Despite these small differences, both trees are largely congruent, and agree on most of the major groups.

A study that focuses on the distribution of thermophilic fungi throughout the tree has been published by Morgenstern et al (68) and we evaluated conflicts and congruence between this published phylogeny and ours. Both phylogenies manifest a perfect match in most groups, except for some small discrepancies in Basidiomycota. Specifically, *Heterobasidion annosum* is placed differently in the two trees: next to A*garicales* by Morganstern *et al*, and on a sister branch in our tree. Apart from this difference, the rest of the topology is similar in both trees.

Next, we proceeded to compare our tree with several other studies by using nodal distance and split distance measurements provided by TOPD/FMTS, which can be seen in Table 1. The first study we used for comparison with our phylogeny and the largest one, focusing on the phylogeny of the Basidiomycota, is a six-gene phylogeny by Zhao et al (69). A Tanglegram representing the comparison of both phylogenies can be seen in Figure ***5***. Both phylogenies agree for most part of the tree, especially in the Agaricomycotina subphylum (see Figure ***5***), which agrees across both trees, except for the placement of *Tricholoma matsutake* and *Heterobasidion annosum*. Nevertheless, disagreements between the structure of the tree can be found in the subphylums of Basidiomycota, Pucciniomycotina and Ustilaginomycotina. The disagreement between both trees comes from the placement of these subphylums, where in our phylogeny Ustilaginomycotina is found as a sister clade to Agaricomycotina, in *Zhao et al* phylogeny Pucciniomycotina is found as the sister clade to Agaricomycotina. The branch split between the Ustilaginomycotina and Agaricomycotina clades in our phylogeny is not robustly supported (58%) and the groupings proposed by Zhao *et al* phylogenetic hypothesis is has additional support from recent literature (70). Moreover, the reduced tree and data heterogeneous phylogeny recovered in our study is also in agreement with the Zhao *et al* hypothesis, as can be observed in Figure ***2***. Therefore, we can assume that the branch split between Pucciniomycotina and Ustilaginomycotina proposed by the complete phylogeny hypothesis is incorrect. Other than this discrepancy, both phylogenies agree on the remaining topology of the tree.

**Figure 5:**
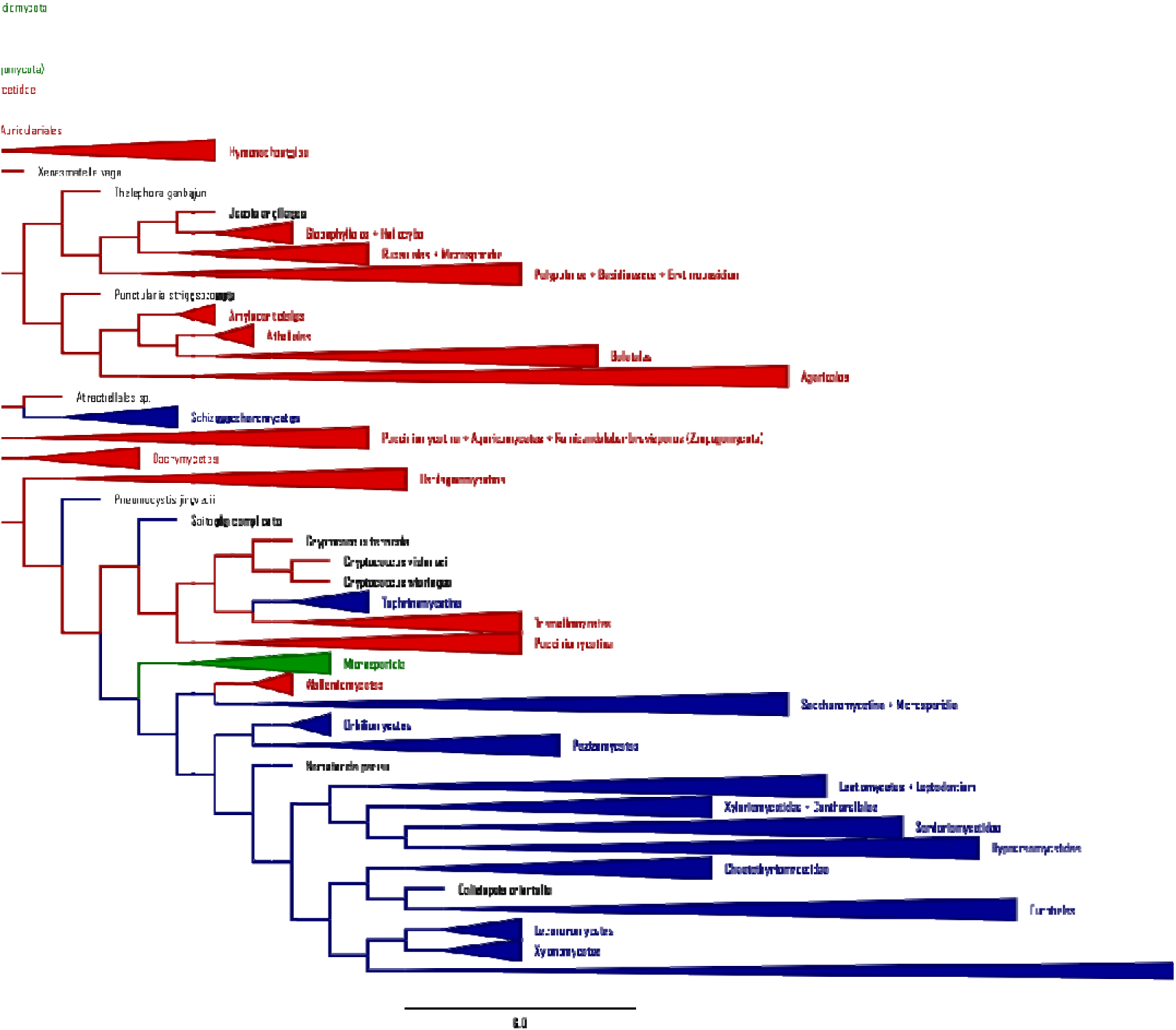
Supertree reconstruction using the MRP method. Branches are collapsed when they are monophyletic according to NCBI’s taxonomy. In Green are the groups of early fungi outside of Dikarya, Red represents Basidiomycota, Blue represents Ascomycota and Black is the out-group, *Rozella allomycis*.

**Figure 5:**
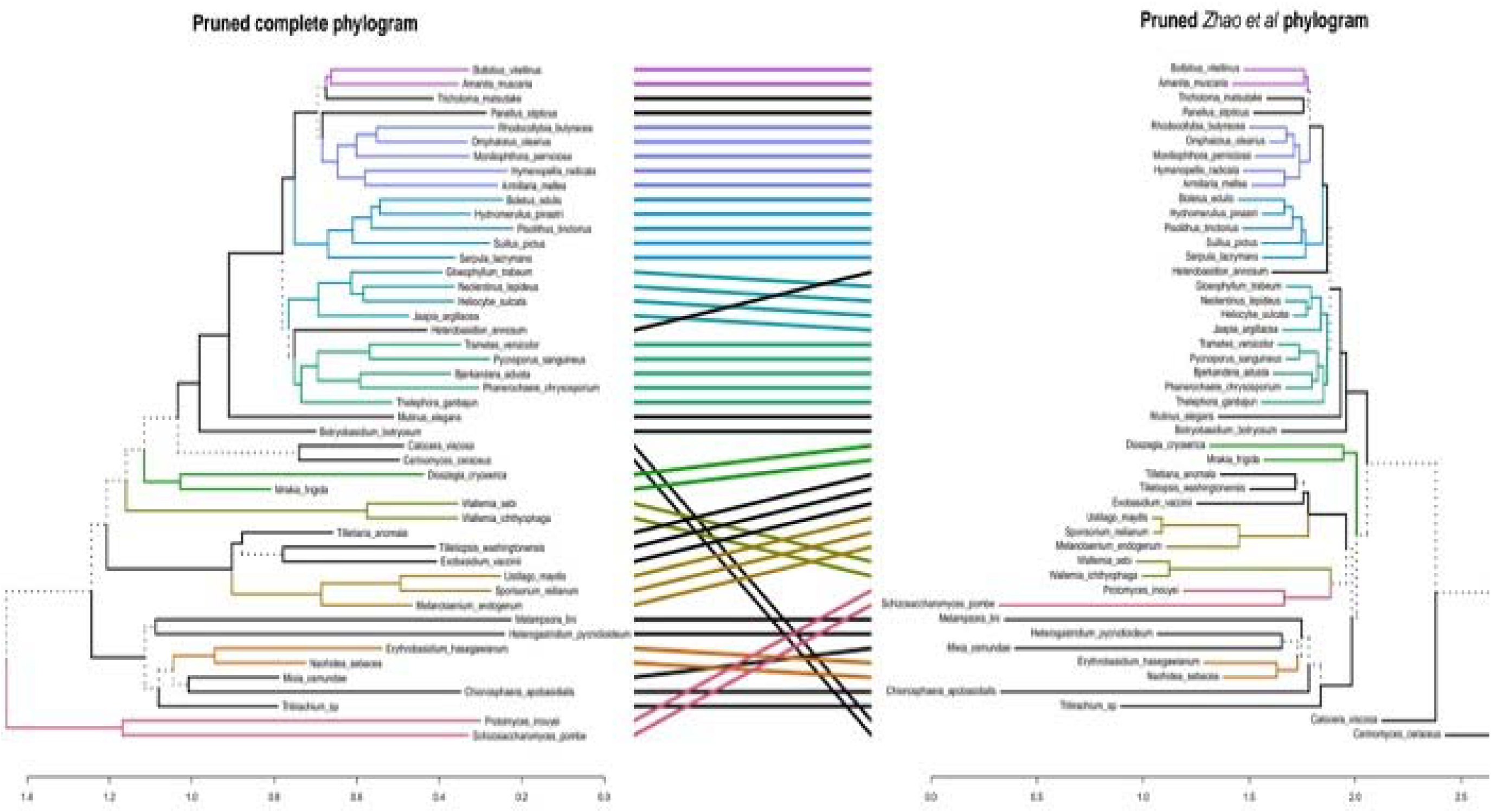
Tanglegram of the complete dataset phylogenetic tree (left) compared with Zhao et al six-gene phylogeny of the Basidiomycota (right). The project tree is focused on the Basidiomycota region. Dotted lines indicate differing branches between the phylogenies. Branches in the phylogenies are assigned colours to facilitate viewing.

We next compared our tree with a phylogeny focused in the Gloeophyllales order in the Agaricomycetes class proposed by Chen *et al* (*63*). The tanglegram representing this comparison can be seen in Figure ***7***. In this case, 16 taxa are shared between both phylogenies, and we observe disagreement in the placement of three of these taxa. The first source of disagreement is found in the placement of two taxa, *Punctularia* and *Trametes,* within the Agaricomycetes class. This disagreement could be explained by the difference in representation in the Gloeophyllales order, which is poorly represented in our dataset and is well represented in the Chen *et al* phylogeny. As for the last disagreeing taxa, *Heterobasidion* (a member of the *incertae sedis* groups within the Agaricomycetes class), can be explained by the opposite phenomenon, where the surrounding clades are better represented in our phylogeny, particularly other *incertae sedis* groups. Lastly, the general structure of both phylogenies is similar despite the disagreeing taxa.

**Figure 6:**
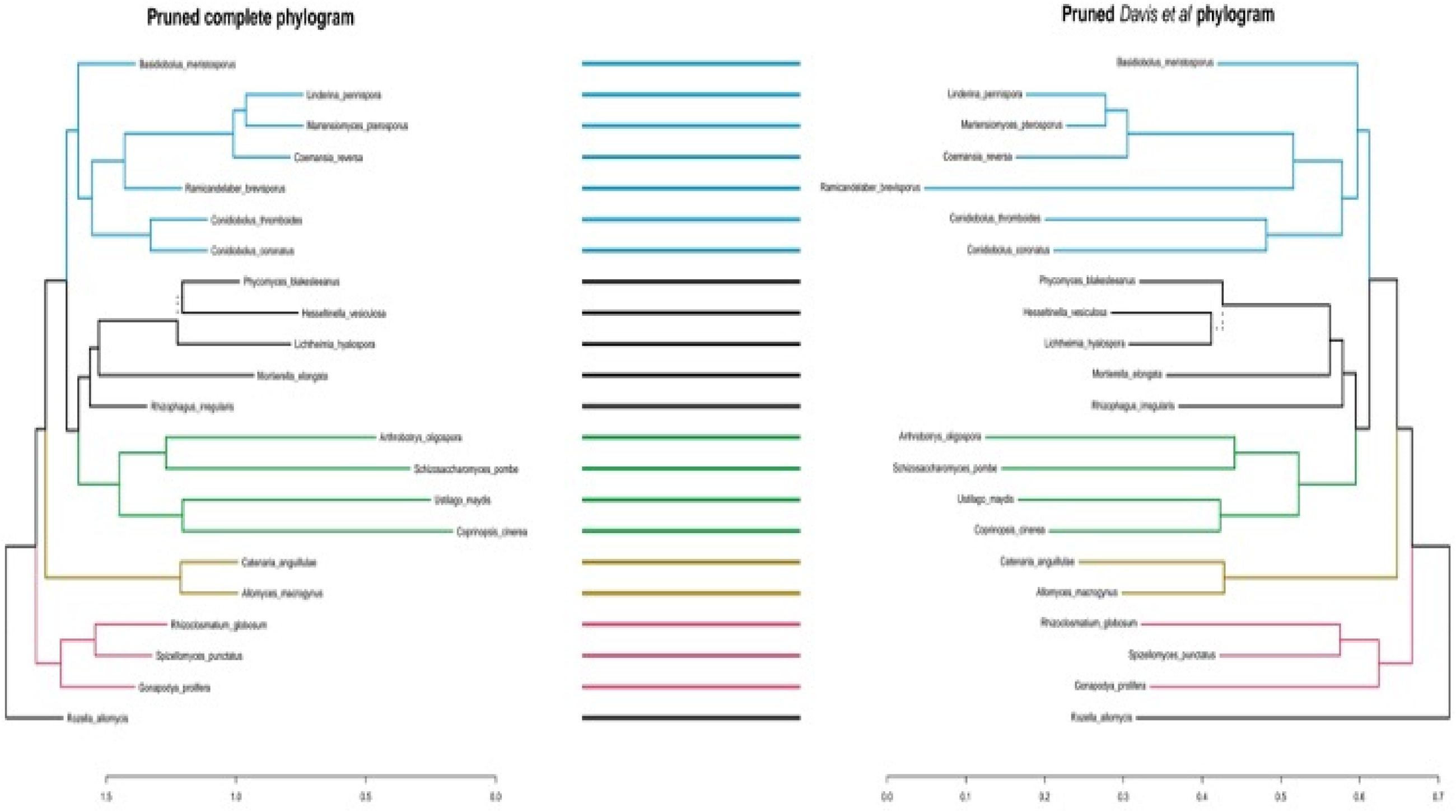
Tanglegram of the complete dataset phylogenetic tree (left) compared with Davis et al phylogeny (right). Dotted lines indicate differing branches between the phylogenies. Branches in the phylogenies are assigned colours to facilitate viewing.

**Figure 7:**
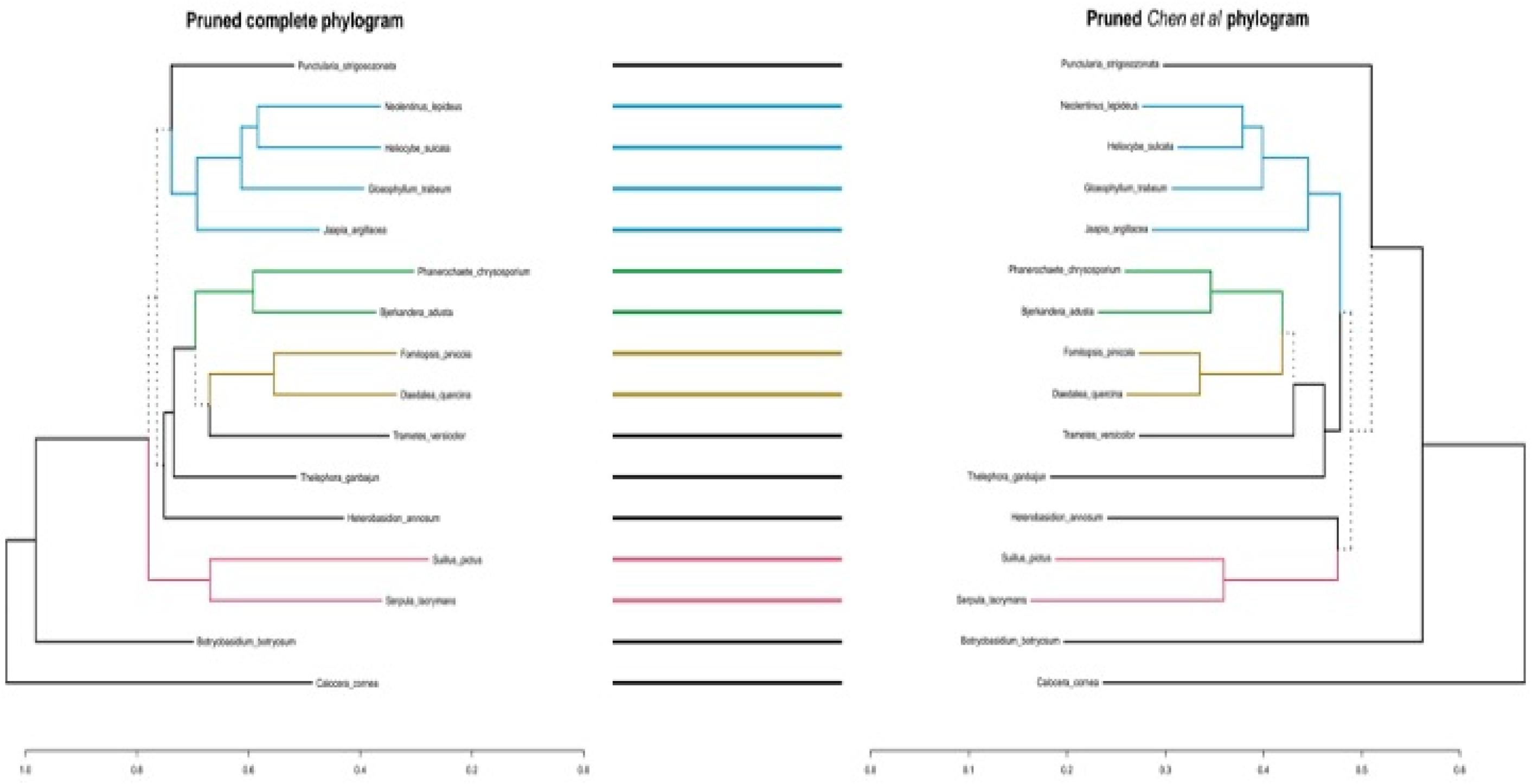
Tanglegram comparing this project phylogenetic tree (left) with Chen et al phylogeny (right). The project tree is focused on the Basidiomycota division. Dotted lines indicate differing branches between the phylogenies. Branches in the phylogenies are assigned colours to facilitate viewing.

The last comparison was carried out with the phylogeny proposed by De Crop *et al* (64), focused on the resolution of the Russulales order and the tanglegram comparing both phylogenies can be seen in Figure ***8***. The disagreements between both phylogenies are of similar nature with Chen *et al* comparison, where there is a difference between the population of the clades where the disagreeing taxa are located. Specifically for this comparison, most of the disagreeing taxa are part of the Mucorales order. As an example, one of the disagreeing taxa is the *Mucor* family, which is represented by three members in our tree and only one member in De Crop *et al* phylogeny. The rest of the disagreeing taxa have similar issues, being part of well represented clades in our phylogeny and comparatively less dense clades in De Crop *et al* phylogeny. Still, both trees nodal and split distances are low, so the general structure of the trees is mostly in agreement.

**Figure 8:**
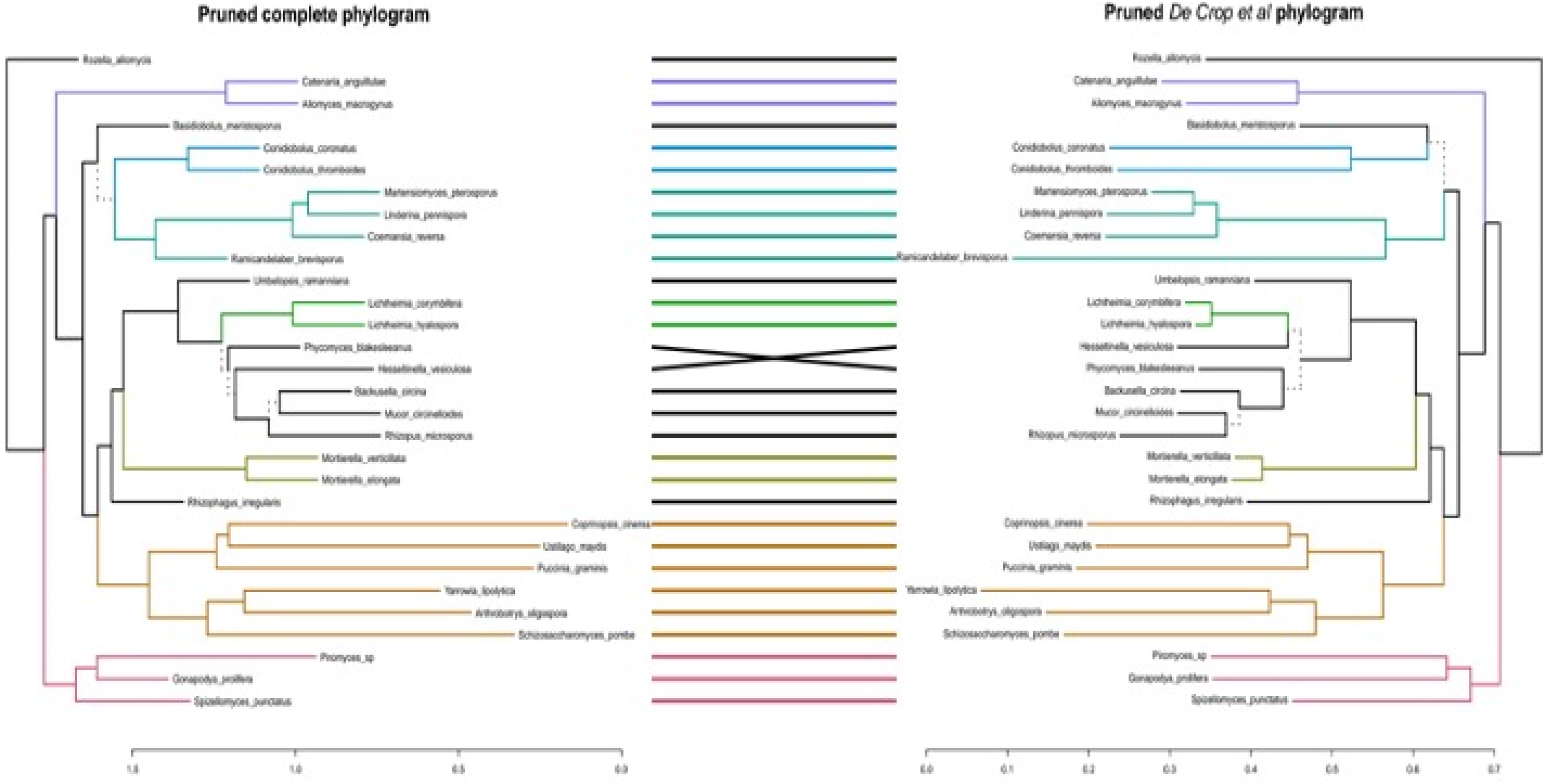
Tanglegram comparing this project phylogenetic tree (left) with De Crop et al phylogeny (right). Dotted lines indicate differing branches between the phylogenies. Branches in the phylogenies are assigned colours to facilitate viewing.

The rest of the phylogenetic comparisons are focused on smaller phylogenies that deal with recovering the placement of specific orders, so they encompass smaller sections of the fungal taxonomy. The next study used for the assessment of our phylogeny is a phylogeny proposed by Davis *et al* (*62*), focused on the resolution of the *Zoopagales* order while using a similar maximum-likelihood approach to the analysis conducted in this project but with a different dataset. The tanglegram comparing both phylogenies can be observed in Figure ***6***. There are disagreements in two out of the 22 taxa that are commonly shared between both phylogenies, and all disagreeing taxa belong to the order *Mucorales*. These two disagreeing taxa show low branch split support in our phylogenetic hypothesis, *Phycomyces blakesleeanus* and *Hesseltinella vesiculosa* (57% and 55% bootstrap support respectively), so it is likely that the branch split in our phylogeny for these species are incorrect. The rest of the taxa and the general structure of both phylogenies are in agreement.

To conclude, our proposed phylogenetic hypotheses are mostly in agreement with previously proposed phylogenies in the literature, though we have recovered several clades with low support that would benefit from a denser sampling. Many of the disagreements between phylogenies could be explained due to the different density of taxa in the clades, but for the most part we have recovered strongly supported clades that are largely in agreement with other studies.

## Models

In the initial steps of our phylogenetic analysis, the simpler maximum likelihood models agreed on the same topology when constructing trees in different runs before the outgroups were included in the alignment. However, when we included outgroups in the analysis the shortcomings of simpler maximum likelihood models became apparent, particularly in relation to the placement taxa whose placement is known to be problematic, especially Microsporidia whose rate of evolution is different when compared with the rest of the tree. The maximum likelihood models would often split the outgroups, not agreeing in any particular topology, and usually resulted in topologies that were poorly supported. Even the more parameter-rich homogeneous maximum likelihood models that account in some way for different rates of evolution in different parts of the tree were not able to recover a well supported phylogeny.

To properly account for phylogenetic signals in our dataset we employed heterogeneous likelihood models. Heterogeneous models can account for variation in evolutionary rates across the tree. Relaxing the constraint that evolution is homogeneous across the tree can sometimes result in significantly higher likelihoods, as well as the recovery of different tree topologies. When we used this approach with our reduced dataset, we were recovered a topology that placed the root reliably in repeated reconstructions of the fungal phylogeny. We used two different models for this purpose, a data heterogeneous model, which agreed with the general topology of maximum likelihood models, and a data-and-tree heterogeneous model.

The data-and-tree heterogeneous model has shown topological discrepancies compared to less parameter-rich models, specifically the placement of Microsporidia as a sister branch of the Dikarya subkingdom, and Zoopagomycota and Mucoromycota being a monophyletic group. We have thoroughly tested all the models and even if the data show that all the phylogenetic hypotheses can plausibly explain the data we observe, the data-and-tree heterogeneous model results in a significantly higher likelihood score than the data heterogeneous model, and both models have significantly higher likelihoods when compared to homogeneous models. This suggests that heterogeneous models are better able to explain the data observed in our dataset. One thing of particular relevance is that in every model except the data-and-tree heterogeneous model, all long branches appear clustered together (in our tree these are the out-groups, *Rozella*, and Microsporidia). Therefore, it seems likely that homogeneous and data heterogeneous models are inappropriate for the resolution of the long branch attraction problem that adding a branch with high substitution rates like Microsporidia brings, and that in our hands, it was only when we used the data-and-tree heterogeneous model we could solve this problem. This opens the possibility that Microsporidia are not early fungi, and they instead share a more recent common ancestor with Dikarya that specialised in intracellular parasitism. Convergent evolution towards intracellular parasitism and genome reduction could also explain why they are so often placed together with *Rozella* and other Cryptomycota, since they are also obligate intracellular parasites.

Furthermore, we explored our methodology by construction of a heterogeneous maximum likelihood tree based on the dataset by McCarthy and Fitzpatrick. Our phylogeny, seen in Figure **4**, shares a similar general structure with McCarthy and Fitzpatrick. Both phylogenies agree with placement of the major groups of the fungi, with groups like Basidiomycota being identical in both phylogenies. There are some minor discrepancies in some smaller groups, *i.e. Magnaporthe grisea* is a sister group of *Phaeoacremonium aleophilum* in their phylogeny, while in our phylogeny *Magnaporthe grisea* is a sister group of *Ophiostoma piceae* and *Phaeoacremonium aleophilum* shares a common ancestor with them. The most obvious difference between our phylogenies is the placement of *Rhizophagus irregularis* and *Gonapodya prolifera*. *Rhizophagus irregularis* is next to Dikarya in McCarthy and Fitzpatrick’s phylogeny, while in our phylogeny it appears within Mucoromycota. *Gonapodya prolifera* appears as a sister branch to Neocallimastigomycota, while in our tree it appears within Chytridomycota. However, the placement of these species is recovered as with our validation phylogeny by some of the other phylogenetic approaches used by McCarthy and Fitzpatrick in their study. Overall, both trees are structurally similar with a few minor differences. The similarity between our validation phylogeny and the McCarthy and Fitzpatrick phylogeny indicates that our phylogenetic methodology can be successfully applied to smaller and less complex datasets of fungi and still recover a similar phylogeny compared with what was already published in the literature.

## Conclusion

By using a combination of data heterogeneous maximum likelihood models and data heterogeneous and data-and-tree heterogeneous Bayesian models we were able to construct a fungal phylogeny which has reliably placed the major groups of the fungi in the phylogeny. Many other small groups were also placed in the phylogeny in accordance with recent literature. Tree and data heterogeneous models have helped us resolve deep phylogenetic trees with very differing substitution rates and opened the possibility that Microsporidia are not early fungi but instead form a monophyletic group together with Dikarya, and that Zoopagomycota and Mucoromycota are also a monophyletic group.

---

**Supplementary Figure 1:**
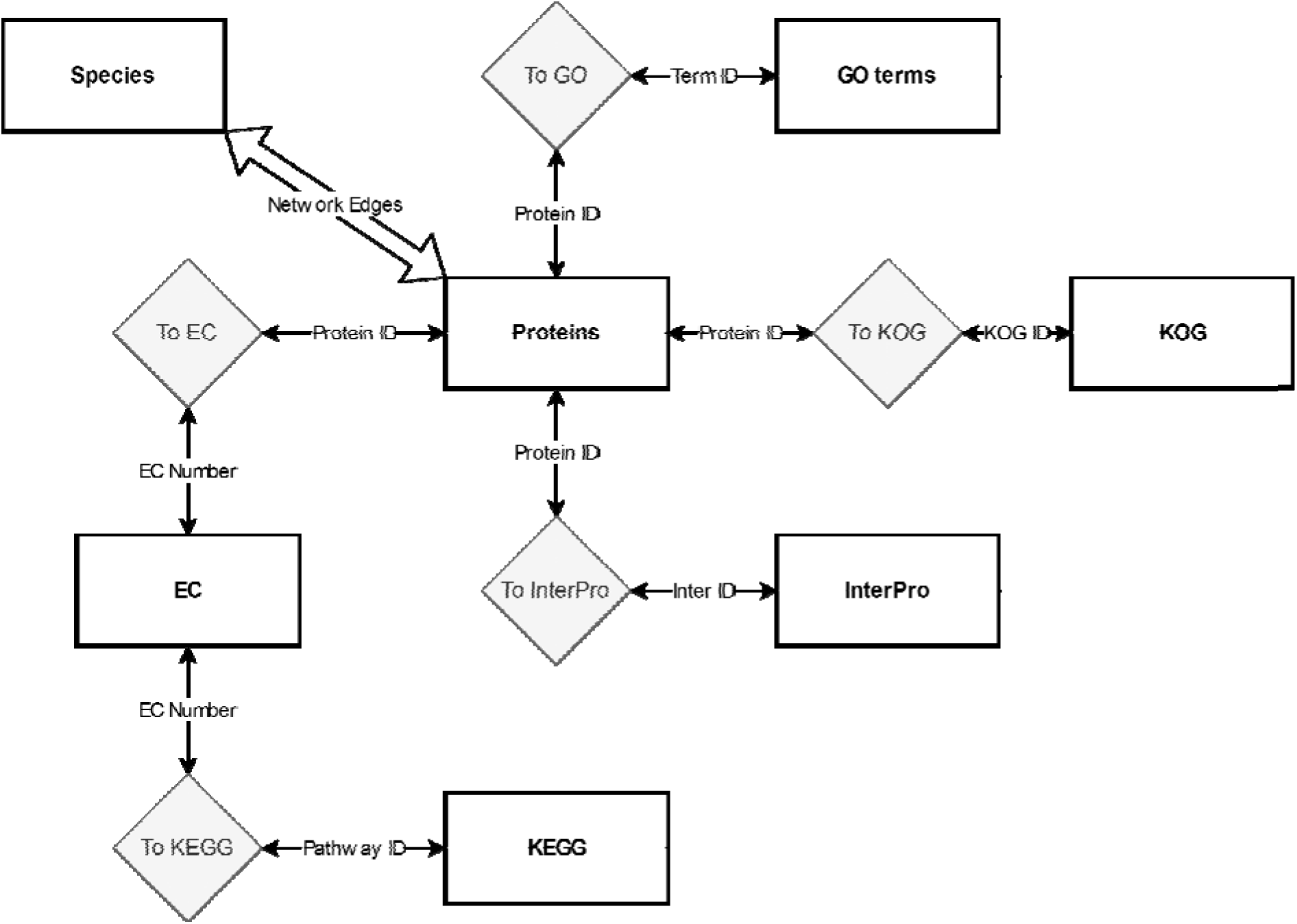
Diagram of the database tables and relationships. Rectangles represents tables that store information and diamonds represent relational tables that store the connections between tables. Each link indicates the identifier used to connect the tables.

